# Rice potassium transporter OsHAK18 mediates phloem K^+^ loading and redistribution

**DOI:** 10.1101/2023.01.02.522451

**Authors:** Like Shen, Qi Wu, Wenxia Fan, Junxia Luan, Na Li, Di Chen, Quanxiang Tian, Wen Jing, Wenhua Zhang

**Affiliations:** State Key Laboratory of Crop Genetics and Germplasm Enhancement, College of Life Sciences, Nanjing Agricultural University, Nanjing 210095, China

**Keywords:** K^+^ transporters, OsHAK18, K^+^ deficiency, phloem loading, rice

## Abstract

High-Affinity K^+^ transporters/K^+^ Uptake Permeases/K^+^ Transporters (HAK/KUP/KT) are important pathways mediating K^+^ transport across cell membrane, which function in maintaining K^+^ homeostasis during plant growth and stress response. An increasing number of studies have shown that HAK/KUP/KT transporters play important roles in potassium uptake and root-to-shoot translocation. However, whether some HAK/KUP/KT transporters mediate K^+^ redistribution in phloem remains unknown. In this study, we revealed that a phloem-localized HAK/KUP/KT transporter, OsHAK18 operated as a typical KUP/HAK/KT transporter mediating cell K^+^ uptake when expressed in yeast, *E. coli* and *Arabidopsis*. It was localized at plasma membrane. Disruption of *OsHAK18* rendered rice seedlings insensitive to low-K^+^ stress. Compared with WT, the *oshak18* mutants accumulated more K^+^ in shoots but less K^+^ in roots, leading to a higher shoot/root ratio of K^+^ per plant. Although disruption of *OsHAK18* doesn’t affect root K^+^ uptake and K^+^ level in xylem sap, it significantly decreases phloem K^+^ concentration and inhibits root-to-shoot-to-root K^+^ translocation in split-root assay. These results reveal that OsHAK18 mediates phloem K^+^ loading and redistribution, whose disruption is favor of shoot K^+^ retention under low-K^+^ stress. Our findings not only reveal a unique function of rice HAK/KUP/KT family member, but also provide a promising strategy to improve rice tolerance under K^+^ deficiency.

## INTRODUCTION

Potassium (K^+^) is one of the most important macroelements and plays crucial roles in regulating plant growth and response to environmental stimuli (Anschütz et al., 2014; Wang and Wu, 2013). As the most abundant cation in plant cells, K^+^ is important for charge neutralization, maintenance of membrane potential, enzyme activation and photosynthesis. K^+^ also functions as a major osmoticum, regulating stomatal movement, phloem transport and cell elongation (Cherel and Gaillard, 2019; Maathuis, 2009; Véry and Sentenac, 2003). Therefore, it is of great significance to maintain K^+^ homeostasis in plants (Anschutz et al., 2014; Srivastava et al., 2020). However, K^+^ deficiency in soil, also called low-K^+^ stress easily results in disruption of plant K^+^ homeostasis and inhibition of growth, development and reproduction (Wang and Wu, 2015).

K^+^ absorption and long-distance translocation are crucial for plants to cope with low-K^+^ stress, which are mediated by many kinds of K^+^ transporters. (Wang et al., 2021c). Therein HAK/KUP/KT transporters play important roles, and are widely found in higher plants (Li et al., 2018; Very et al., 2014). In Arabidopsis, there are 11 members in HAK/KUP/KT family, among which the AtHAK5 has been the most intensively studied object. AtHAK5 is a major pathway of root K^+^ uptake when external K^+^ is below 100 μM (Pyo et al., 2010; Rubio et al., 2010). AtHAK5 is regulated at transcription level and post-translation level. Low-K^+^ stress dramatically induced the expression of AtHAK5, which was dependent on the ethylene-elicited ROS production (Jung et al., 2009; Shin and Schachtman, 2004; Wang et al., 2021a). A transcription factor, ARF2 represses AtHAK5 expression under normal condition, and can be phosphorylated for degradation in response to low-K^+^ stress (Zhao et al., 2016). RAP2.11 and MYB77 can directly bind with HAK5 promoter and facilitate its expression (Feng et al., 2021; Kim et al., 2012). In addition, AtHAK5 can be phosphorylated by CIPK23 to reach maximal transport activity (Ragel et al., 2015; Rodenas et al., 2021). The AtHAK5’s function is conserved in higher plants, its orthologs had been identified in a great number of species, such as rice (Chen et al., 2015), maize (Qin et al., 2019), tomato (Nieves-Cordones et al., 2020), etc. Besides AtHAK5, some other HAK/KUP/KT transporters’ roles were also reported. AtKUP7 is Involved in K^+^ acquisition and root-to-shoot translocation under low-K^+^ stress (Han et al., 2016). Three members (AtKUP2,6,8) mediate K^+^ efflux and play key roles in osmotic adjustment by balancing K^+^ homeostasis in cell growth and drought stress responses (Osakabe et al., 2013). AtKUP9 is located at ER membrane, and mediates K^+^ and auxin release from ER in root quiescent center (QC) cells to maintain meristem activity (Zhang et al., 2020).

In rice, HAK/KUP/KT family contains 27 members, divided into four subgroups (Gupta et al., 2008; Very et al., 2014). OsHAK1, 5, 8 and 16 are involved in root K^+^ uptake and translocation from roots to shoots, and also play important roles in rice salt tolerance (Chen et al., 2015; Feng et al., 2019; Horie et al., 2011b; Wang et al., 2021b; Yang et al., 2014). OsHAK21also contributes to maintenance of K^+^ homeostasis as a K^+^ transporter under salt stress, whose expression was induced by NaCl but not K^+^ deficiency (He et al., 2019; Shen et al., 2015). OsHAK12 was shown to be permeable to Na^+^ but not K^+^, and possibly retrieved NaCl from xylem vessel, thereby reducing NaCl content in shoots (Zhang et al., 2021).

An increasing number of studies have shown that HAK/KUP/KT transporters play important roles in potassium uptake and root-to-shoot translocation (Li et al., 2018; Ragel et al., 2019), which are two tightly coordinated processes (Nieves-Cordones et al., 2019a). However, whether some HAK/KUP/KT transporters mediate K^+^ redistribution in phloem tissue remains unknown. In this study, we revealed that a phloem-localized HAK/KUP/KT transporter, OsHAK18 operated as a typical KUP/HAK/KT transporter mediating cell K^+^ uptake when expressed in yeast, *E. coli* and *Arabidopsis*. It was localized at plasma membrane. Disruption of *OsHAK18* rendered rice seedlings insensitive to low-K^+^ stress. Compared with WT, the *oshak18* mutants accumulated more K^+^ in shoots but less K^+^ in roots, leading to a higher shoot/root ratio of K^+^ per plant. Although disruption of *OsHAK18* doesn’t affect root K^+^ uptake and K^+^ level in xylem sap, it significantly decreases phloem K^+^ concentration and inhibits root-to-shoot-to-root K^+^ translocation in split-root assay. These results reveal that OsHAK18 mediates phloem K^+^ loading and redistribution, whose disruption is favor of shoot K^+^ retention under low-K^+^ stress. Our findings not only reveal a unique function of rice HAK/KUP/KT family member, but also provide a promising strategy to improve rice tolerance under K^+^ deficiency.

## RESULTS

### OsHAK18 mediates cell K^+^ absorption when expressed in yeast, *E. coli* and *Arabidopsis*

To investigate the transporter activity of OsHAK18, we expressed OsHAK18 in K^+^ uptake-deficient yeast strain CY162, and examined the effect of OsHAK18 expression on the yeast growth. As shown in Fig. 1a, the CY162 transformed with pYES2 empty vector was unable to grown on the low-K^+^ medium containing 0.5 or 1 mM K^+^, but OsHAK18 expression can rescue this growth defect. There were no significant differences among all the transformants when they were grown on AP medium with sufficient K^+^ (10 mM). We also found that the No. 2# transformant expressing *OsHAK18* showed higher capability to grow on LK medium than the clone 1# (Fig. 1a), possibly because the clone 2# has a higher expression level of *OsHAK18* (Fig. 1b).

**Figure 1.**
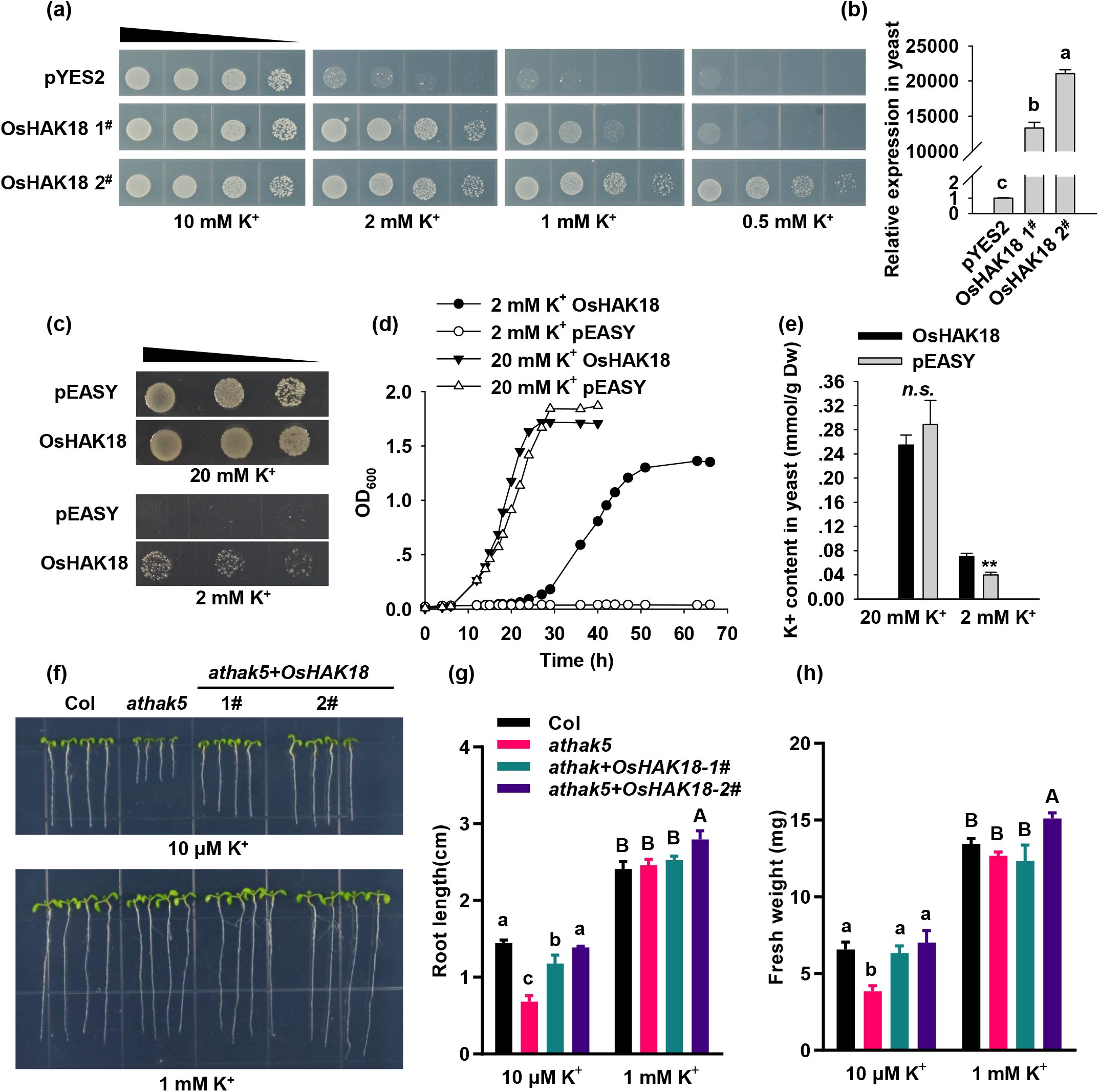
Functional characterization of OsHAK18 in K^+^ uptake-deficient yeast (CY162), *Escherichia coli* (TK2420), and *Arabidopsis athak5*. (a) Growth status of K^+^ uptake-deficient yeast (CY162) expressing *OsHAK18* and empty vector (pYES2) on arginine phosphate (AP) medium supplemented with different concentrations of KCl (0.5, 1, 2, 10 mM). Black triangle indicates 1:10 serial dilutions of yeast cells placed on the mediums. Two independent clones of *OsHAK18* transformants (1^#^ and 2^#^) were tested. (b) Expression level of *OsHAK18* in the transformants assessed by qRT-PCR. The PCR templates were cDNA reverse-transcripted from yeast RNA. The yeast housekeeping gene *ACT1* was used as an internal control. (c) Growth status of in K^+^ uptake-deficient *E. coli* (TK2420) expressing *OsHAK18* and empty vector (pEASY) on control medium (containing 20 mM KCl) or LK medium (containing 2 mM KCl).). (d) The cell growth curve of the TK2420 transformants under control and LK conditions. (e) K^+^ contents in yeast cells. Dw, dry weight of yeast. (f) Expression of *OsHAK18* in *Arabidopsis athak5* mutant relieved its low-K+-sensitive phenotype. The wild type (Col), *athak5* and OsHAK18-expressed *athak5* (athak5+OsHAK18) plants were grown for 8 d on medium containing KCl at the concentrations indicated. (g,f) Root length and fresh weight of the 8-day-old plants. Letters denote significantly different groups identified by Tukey’s multiple comparison test (p < 0.05). *n*.*s*. indicates non-substantial differences at that level of significance

To further verify the K^+^ transport function of OsHAK18, we performed complementation tests using the K^+^ uptake-deficient *E. coli* strain TK2420. The transformants expressing *OsHAK18* can grow on the relative low K^+^ medium, whereas the transformants harboring empty vector cannot (Fig. 1c), which was further confirmed by the growth curve analysis (Fig. 1d). In addition, under low K^+^ condition, the transformants expressing *OsHAK18* accumulated more K^+^ than the empty vector control (Fig. 1e). These findings are consistent to the result of yeast growth assay, indicating that OsHAK18 can confer K^+^ uptake capability to yeast and *E*.*coli*.

To further characterize the transport activity of OsHAK18 in plant cells, we expressed *OsHAK18* in *Arabidopsis athak5* mutant, which displayed a low-K^+^-sensitive phenotype (Pyo et al., 2010). The ectopic expression of OsHAK18 in *athak5* was driven by a 2×35S promoter. Thus, the phenotypic complementation of *athak5* by OsHAK18 may be observed if OsHAK18 also operated as a K^+^ transporter like AtHAK5. Two independent homozygous transgenic lines (*athak5*+*OsHAK18*, 1# and 2#) were obtained (Fig. S1) and used in LK phenotype analysis. As shown in Fig. 1f, the *athak5* mutants showed severe growth defects on LK medium (5 μM K^+^), as exemplified by shorter root length and less fresh weight (Fig. 1g,h). However, these low-K^+^-sensitive phenotypes were almost completely suppressed in the two *OsHAK18*-expressed transgenic lines (*athak5*+*OsHAK18*, 1# and 2#). There were no significant phenotypic differences among all the genotypes grown on the control medium containing 1 mM K^+^ (Fig. 1f). The root length and fresh weight of the *OsHAK18* transgenic line 2# were significantly higher than the corresponding values of the line 1# and WT under control condition, possibly due to the higher expression level of *OsHAK18* in the line 2# (Fig. S1).

Considering that some KUP/HAK/KT transporters were shown to be permeable to Na^+^ (Zhang et al., 2021; Zhang et al., 2019), we also determined whether OsHAK18 mediated cell Na^+^ absorption using the Na^+^-sensitive yeast B31. AtHKT1;1, which mediates Na^+^ uptake, was used as a positive control. With the increase of NaCl concentration in medium, the growth of all the B31 transformants was gradually inhibited. However, AtHKT1;1-expressing cells showed a hypersensitive phenotype to Na^+^ treatment, whereas the performance of OsHAK18-expressing cells was similar to that of the empty vector control (Fig. S2), suggesting that OsHAK18 may be not permeable to Na^+^.

### Subcellular localization and expression pattern of OsHAK18

To investigate the subcellular localization of OsHAK18, We expressed OsHAK21 fused to EGFP at the C-terminal (OsHAK18:EGFP) in onion epidermal cells. As shown in Fig. 2a, OsHAK18-EGFP was confined to the cell periphery. Plasmolysis assays further suggested that OsHAK18 was localized at the plasma membrane. To confirm the plasma membrane localization of OsHAK18, we also expressed OsHAK18 in *Nicotiana benthamiana* mesophyll cells and used OsAKT1:mCherry as a plasma membrane marker (Li et al., 2014). When OsHAK18:EGFP and OsAKT1:mCherry were co-expressed, the two fluorescence signals overlapped considerably (Fig. 2b). These findings indicate that OsHAK18 is localized at plasma membrane.

**Figure 2.**
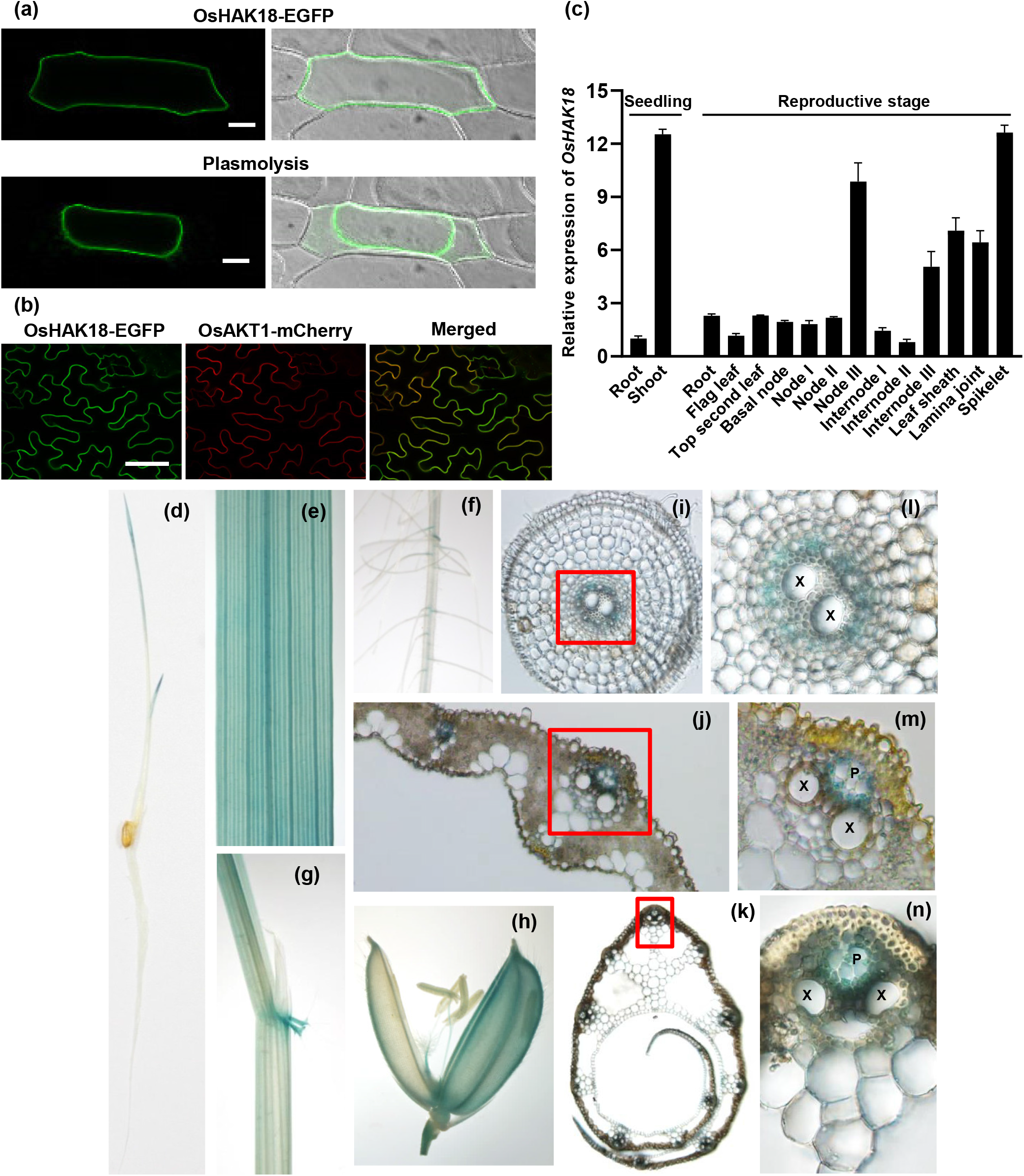
Subcellular localization and expression pattern of OsHAK18. (a) Subcellular localization analysis of OsHAK18-EGFP fusion protein in onion epidermal cells. A plasmolyzed cell expressing OsHAK18-EGFP is shown on the right panel. Bars = 50 μm. (b) Co-localization analysis of OsHAK18-EGFP and OsAKT1-mCherry in tobacco leaves. Representative laser-scanning images of tobacco mesophyll cells co-expressing OsHAK18-EGFP and OsAKT1-mCherry are shown. OsAKT1-mCherry was used as a marker of plasma membrane. Bar = 50 μm. (c) Expression analysis of *OsHAK18* in diverse tissues of rice plants. The relative transcript levels of *OsHAK18* were determined by qRT-PCR and were normalized to the value in seedling roots which was set as 1. (d to n) Tissue-specific expression of *OsHAK18* was analyzed by using the β-glucuronidase (GUS) reporter driven by *OsHAK18* promoter. (d) A 5-day-old seedling. (e to g) GUS signal was observed in leaf, root, lamina joint, leaf auricles and sheath of a 21-day-old seedling. (h) Glumous flower. Transverse cross sections of root (i), leaf blade (j) and leaf sheath (k) were shown. (l), (m) and (n) are enlargements of the red boxes in (i), (j) and (k), respectively.

RT-PCR analysis showed that OsHAK18 was more preferentially expressed in shoots than in roots during rice seedling stage, while it was highly expressed in spikelet, Node III, leaf sheath and etc. during reproductive stage (Fig. 2c). To profile the expression pattern of OsHAK18 in more detail, we generated transgenic plants carrying GUS reporter gene driven by *OsHAK18* promoter, and perform GUS staining. In a 5-day-old seedling, strong GUS signals can be only detected in leaves (Fig. 2d). However, in a 21-day-old seeding, leaves and roots were dyed, especially in vascular tissues (Fig. 2e,f,). GUS signal were also shown in leaf auricles, leaf sheath, pistil, stamen filaments and glumes (Fig. 2g,h). Transverse cross section of root showed that the dyed cells were restricted to the regions which were probably protophloem between points of protoxylem (Fig. 2i,l). In transverse cross sections of leaf blade and sheath, the GUS signals were particularly detected in the phloem tissue, especially in companion cells and phloem parenchyma cells (Fig. 2j,k,m,n).

### Disruption of *OsHAK18* inhibits K^+^ deficiency-induced wilting and chlorosis of leaves

To reveal OsHAK18’s physiological function *in vivo*, we obtained a *Tos17* insertion line (Accession No.: NE5666) from the rice *Tos17* mutant database. The *Tos17* insertion was located in the last exon of the *OsHAK18* gene (Fig. 3a), which was confirmed by PCR analysis using specified primer sets (Fig. S3a, Table S1). We isolated homozygous individuals and verified that the *Tos17* insertion dramatically decreased *OsHAK18* expression level (Fig. S3b). Thus, this homozygous mutant was named *oshak18*. We also generated knockout transgenic lines of *OsHAK18* using the CRISPR/Cas9 editing system. Two specific gRNA target sequence (Target 1 and 2) were designed using CRISPR-PLANT database (Xie et al., 2014). In the T2 generation, we isolated two independent CRISPR mutant lines, which were named *cr-oshak18-1* and *cr-oshak18-2*. As shown in Fig. 3a and Fig. S3c, *cr-oshak18-1* was a biallelic mutant line carrying a single-base insertion (A or T) at the same position in Target 2, while *cr-oshak18-2* was a homozygous mutant with a deletion of 43-bp fragment. Both mutations led to frameshift and premature termination.

**Figure 3.**
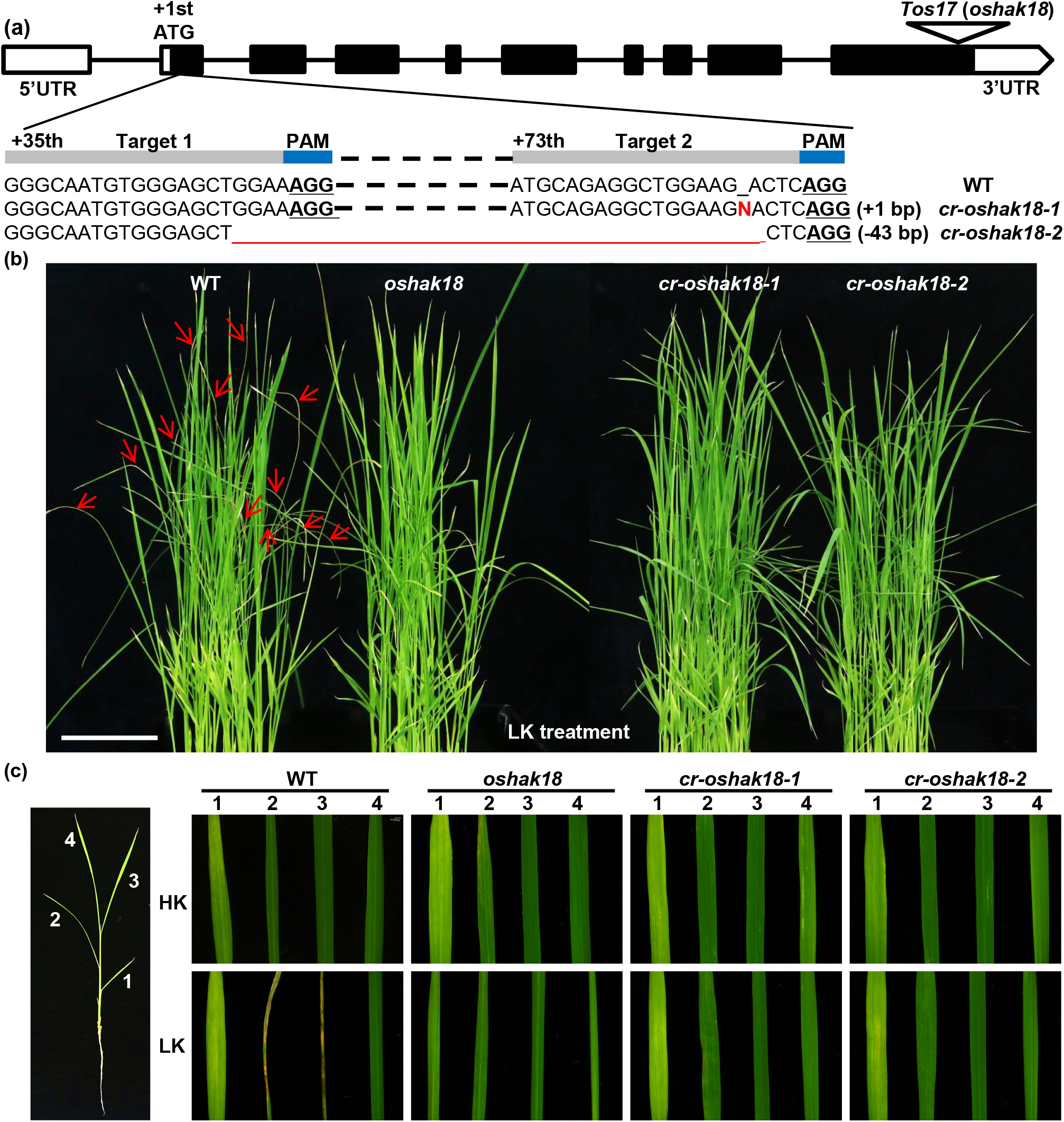
Phenotypic comparison between wild type and *oshak18* mutants under K^+^ deficiency. The *oshak18* mutants were generated using *Tos17* insertion or CRISPR/Cas9 gene editing system. (a) A schematic diagram shows the *Tos17* insertion and the target sites of *oshak18*-sgRNA. Black boxes represent exons, while white boxes indicate the 5’ and 3’ untranslated regions (UTR). Introns are shown by black lines. The triangle on the last exon indicates the *Tos17* insertion position. The triangle at the first exon indicates the target sites and the mutations caused by CRISPR/Cas9 editing in two independent mutant lines (*cr-oshak18-1/2*). The PAM sequence is underlined and bolded. The red letter and the red line in the sequences of the CRISPR lines respectively represent the insertion or deletion relative to WT sequence. N represents A or T (b) Disruption of *OsHAK18* in rice inhibits the K^+^ deficiency-induced wilting and chlorosis of leaves. Two-week-old seedlings were transferred to low-K^+^ (LK) hydroponic medium containing 10 μM K^+^, and treated for 12 days. The red arrows show the wilted chlorotic leaves only observed in WT. Similar results were obtained in three independent experiments. Bars = 5 cm. (c) Representative images show the leaves of WT and *oshak18* mutants grown under control (5 mM) or LK (10 μM) conditions. The numbers (1 to 4) represent four leaves numbered in order of development, which were excised from one seedling of the indicated genotype. A schematic diagram on the right panel shows the numeration and position of leaves.

Given the K^+^ transport activity of OsHAK18 characterized in yeast, *E. coli* and Arabidopsis (Fig. 1), we supposed that OsHAK18 was involved in K^+^ transport in rice and performed phenotypic analysis under K^+^ deficiency condition (low-K^+^ treatment). Two-week-old seedlings of WT, *oshak18* and two CRISPR mutant lines were transferred to low-K^+^ (LK) hydroponic medium containing 10 μM K^+^. After 12-d treatment, the WT seedlings displayed obvious K^+^ deficiency symptom that quite a few leaves became wilted and chlorotic (red arrows in Fig. 3b), whereas the leaves of *oshak18* and two CRISPR lines had no similar symptom (Fig. 3b). There were no significant differences on seedling leaves among these genotypes before LK treatment (Fig. S4).

To compare the phenotype of leaves in more detail, all of the expanded leaves were excised from seedlings and were observed with stereo microscope. The 28-d-old seedlings have four expanded leaves, which were numbered from 1 to 4 in order of development (Fig. 3c). Under high K^+^ (HK) condition, there were no marked differences on each corresponding leaf among WT, *oshak18* and two CRISPR lines. However, after K^+^ deficiency treatment, the 2th and 3rd leaves of WT showed severe wilting and chlorosis, whereas the corresponding leaves of *oshak18* and two CRISPR lines remained unwilted and green (Fig. 3c). The measurements of width and length of leaves showed that there were no differences on length of leaves between WT and *oshak18* regardless of LK treatment, except that the 4th leaves of *oshak18* were shorter than those of WT after LK treatment (Fig. S5). However, the 2nd and 3rd leaves of WT were significant narrower than those of *oshak18* after LK treatment, whereas there were no differences on width of leaves between WT and *oshak18* under HK control condition (Fig. S5). These findings indicated that disruption of *OsHAK18* inhibited K^+^ deficiency-induced wilting and chlorosis of leaves, generating a LK-resistant phenotype in *oshak18* mutants.

### *oshak18* mutants accumulate more K^+^ in shoots but less K^+^ in roots than WT under K^+^ deficiency

To further elucidate the OsHAK18’s function, we determined K^+^ contents in shoots and roots of diverse plant materials, including WT, *oshak18* and the CRISPR line (*cr-oshak18-1*). As shown in Fig. 4a,b, under control condition, no differences were observed among the three genotypes. After LK treatment, K^+^ contents in shoots and roots of all the genotypes were significantly decreased. However, compared with WT, the *oshak18* and *cr-oshak18-1* accumulated more K^+^ in shoots but less K^+^ in roots (Fig. 4a,b). This result suggested that OsHAK18 might affect K^+^ distribution between shoots and roots. Thus, we further quantified the shoot/root ratios of K^+^ per plant. As shown in Fig. 4c, the shoot/root ratios of K^+^ in *oshak18* and *cr-oshak18-1* were markedly higher than that in WT after LK treatment, whereas no differences were observed under HK control condition.

**Figure 4.**
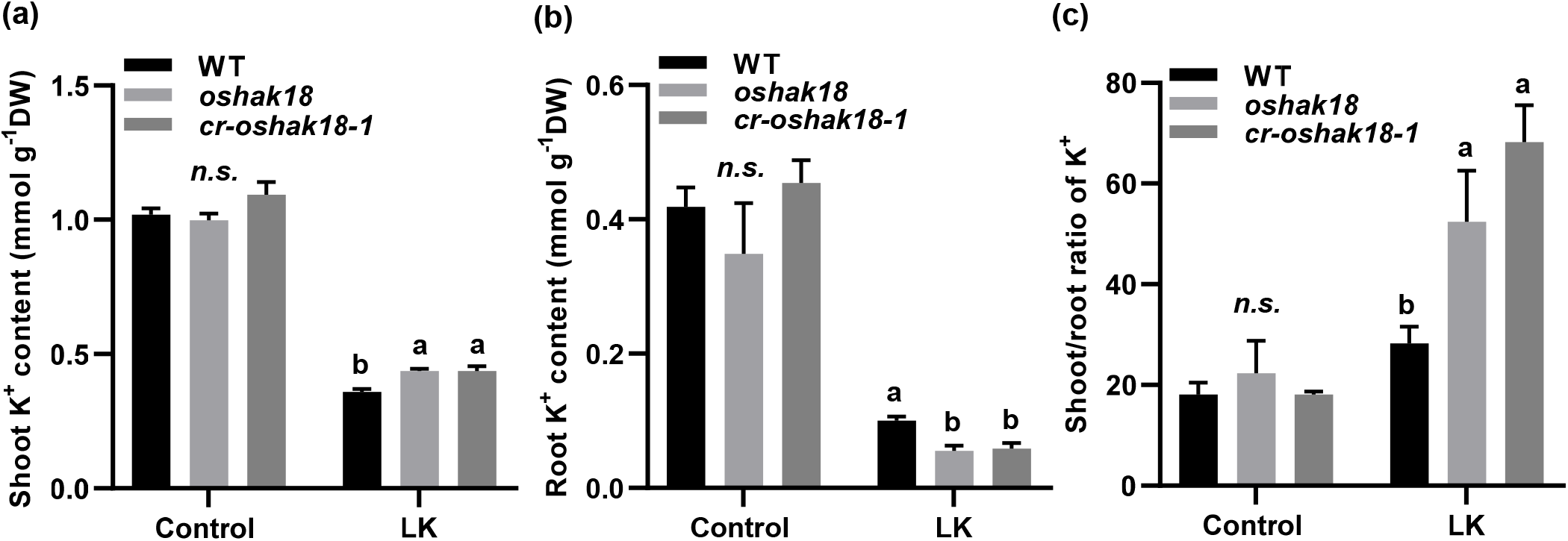
*oshak18* mutants accumulate more K^+^ in shoots but less K^+^ in roots than WT under K^+^ deficiency. Fourteen-day-old rice seedlings were transferred to control (5 mM K^+^) or LK hydroponic medium (10 μM K^+^), and were treated for 14 days. (a) K^+^ contents in shoots (b) K^+^ contents in roots. (c) The shoot/root ratios of K^+^ per plant. Each data represents means ± SD (n = 4). Statistical significance analyzes were carried out by one-way analysis of variance (ANOVA) with Tukey’s multiple range test at P < 0.05. *n*.*s*. indicates non-substantial differences at that level of significance.

To analyze the OsHAK18-regulated K^+^ distribution in shoots in more detail, we further determined the K^+^ contents in each leaf blade and sheath after LK treatment. As shown in Fig. 5a, after LK treatment, the K^+^ contents in the leaf blades (especially the 2nd, 3rd and 4th blades) of *oshak18* mutant lines were significantly higher than the corresponding values of WT, whereas no differences or markedly less differences were found under HK control condition (Fig. 5b). Similar phenomenon was observed in the K^+^ contents analysis on leaf sheath (Fig. 5c,d).

**Figure 5.**
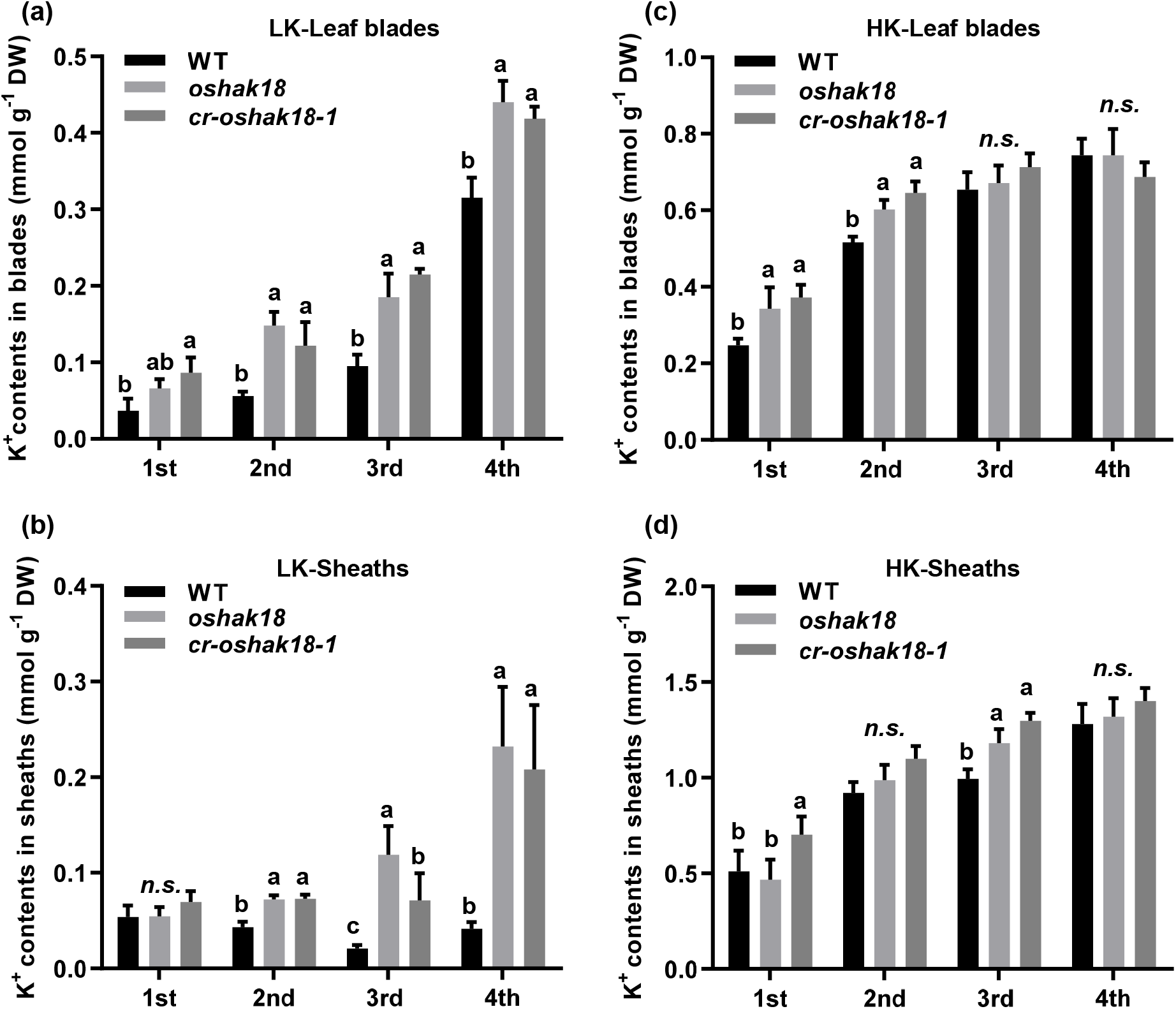
Disruption of *OsHAK18* results in accumulation of K^+^ in leaf blades and sheaths under LK treatment. Two-week-old plants were transferred to control (5 mM K^+^) or LK hydroponic medium (10 μM K^+^), and treated for 14 days. At the end of treatment, the two-week-old plants had five fully-expanded leaves with sheaths and one expanding young leaf, which were labeled from 1st to 4th in order of development. The leaf blades and sheaths of each leaves were separated and collected for K^+^ content analysis. (a,b) K^+^ contents in each leaf blades under control or LK condition. (c,d) K^+^ contents in each leaf sheaths under control or LK condition. Each data bar represents means ± SD (n = 4). Statistical significance analyzes were carried out by one-way analysis of variance (ANOVA) with Tukey’s multiple range test at P < 0.05.

In order to further confirm the difference on K^+^ distribution between WT and *oshak18* mutants, we determined K^+^ level in each leaf and roots using micro-X ray fluorescence (μ-XRF) analysis. Under control condition, the K^+^ level in leaves (or roots) showed no significant differences among the three genotypes (WT, *oshak18* and *cr-oshak18-2*) (Fig. 6a,c). However, after 12-d LK treatment, compared with WT, *oshak18* and *cr-oshak18-2* accumulated significantly more K^+^ in leaves, especially in the 2nd, 4th leaves and the adaxial regions of the 3rd leaves, whereas some roots (longest root branches) of WT had a higher level of K^+^ than those of *oshak18* mutants (Fig. 6b,d). These findings were in line with the results of the K^+^ content determination, suggesting OsHAK18 may play a role in in K^+^ shoot-to-root translocation under K^+^ deficiency.

**Figure 6.**
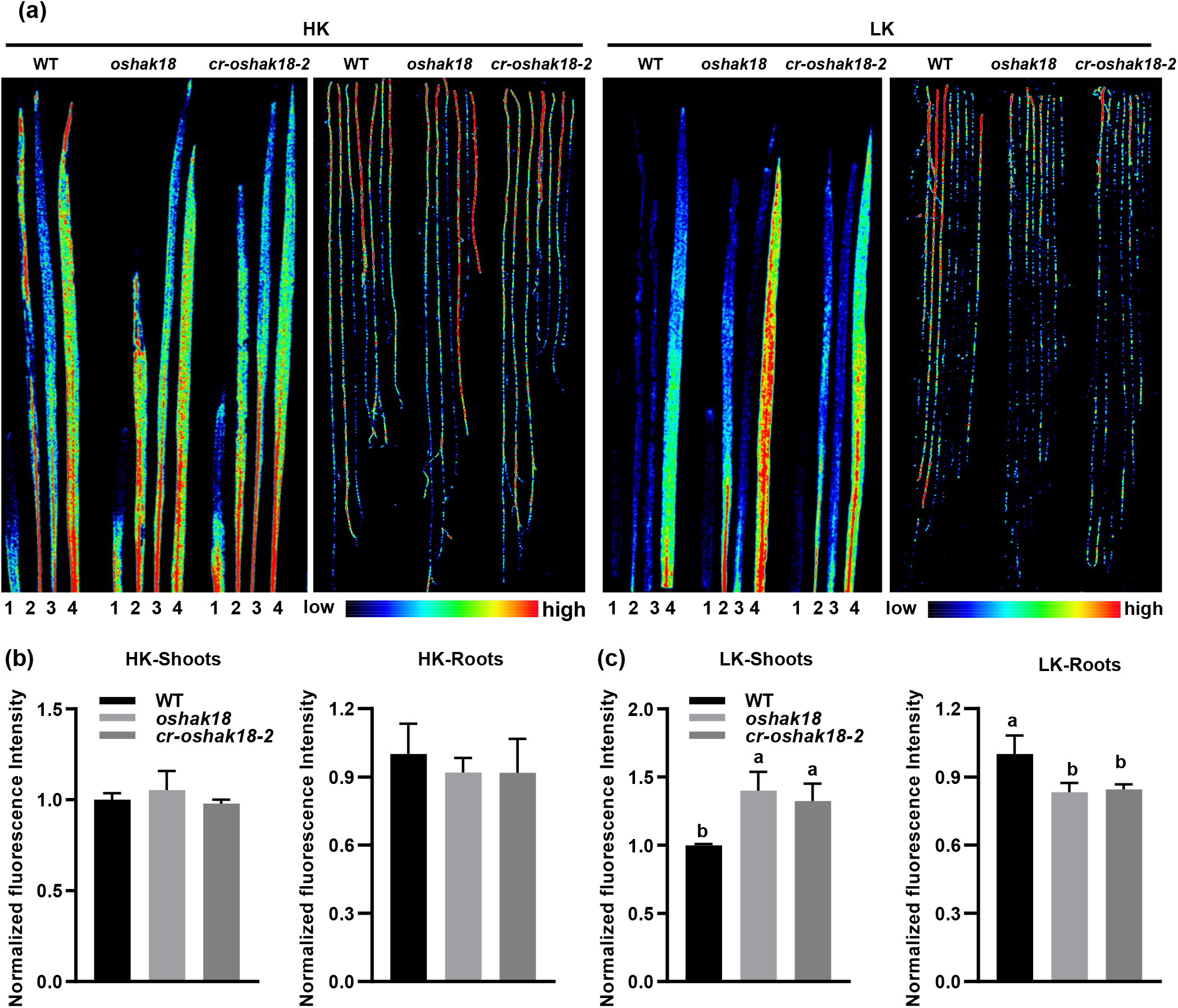
Micro X-ray fluorescence (μXRF) mapping of K^+^ in leaves and roots. Fourteen-day-old rice seedlings were transferred to control (5 mM K^+^) or LK hydroponic medium (10 μM K^+^), and were treated for 12 days. (a) Representative imaging of K^+^ in different genotypes including WT, *oshak18* and *cr-oshak18-2*. The leaves and roots were excised from seedlings and scanned using a μXRF spectrometer. The leaves were arranged from left to right in order of development, labeled from 1 to 4. The relative K^+^ levels in each image are severally represented according to the pseudocolour scale bar. Fluorescence intensity of K^+^ in leaves and roots under HK condition (b) or LK condition (c) was normalized to the corresponding value of WT. Each data bar represents means ± SD (n = 3). Statistical significance analyzes were carried out by one-way analysis of variance (ANOVA) with Tukey’s multiple range test at P < 0.05.

We also generated *OsHAK18* overexpression lines and determined their K^+^ contents. The qRT-PCR analysis showed that the expression level of *OsHAK18* in the two independent lines (OE1 and OE2) was increased to 2.5 or 6.5 times as compared with that in WT (Fig. S6a). These two overexpression lines accumulated more K^+^ in shoots than WT regardless of LK treatment (Fig. S6b). In roots, under control condition, no differences were observed between WT and the overexpression lines, whereas OE2 with a higher expression level of *OsHAK18* had a higher K^+^ content than WT and OE1 after LK treatment (Fig. S6c). The increased K^+^ content in the overexpression lines may be due to the ectopic expression of *OsHAK18* in roots, which may improve the root K^+^ uptake and K^+^ accumulation.

### Disruption of *OsHAK18* doesn’t affect K^+^ uptake and xylem loading but inhibits phloem loading of K^+^

On basis of the results above, we revealed that disruption of *OsHAK18* affected K^+^ distribution between shoots and roots, but whether OsHAK18 contributes to root K^+^ acquisition remains unknown. OsHAK5 had been demonstrated to play an important role in rice root K^+^ acquisition in previous study (Yang et al., 2014). Using K^+^ deletion assay described by Li et al. (2014), we also found that a CRISPR mutant of *OsHAK5* with a Dongjin background showed an obvious defect on K^+^ uptake activity compared with the Dongjin wild type (Fig. 7a), consistent to the finding in previous study. However, the K^+^ deletion curves of *oshak18* coincided almost exactly with that of WT (Nipponbare background) (Fig. 7b), indicating that there was no significant difference on root K^+^ uptake ability between *oshak18* and WT. This notion was further confirmed by the determination of Rb^+^ uptake rate as shown in Fig.7c.

**Figure 7.**
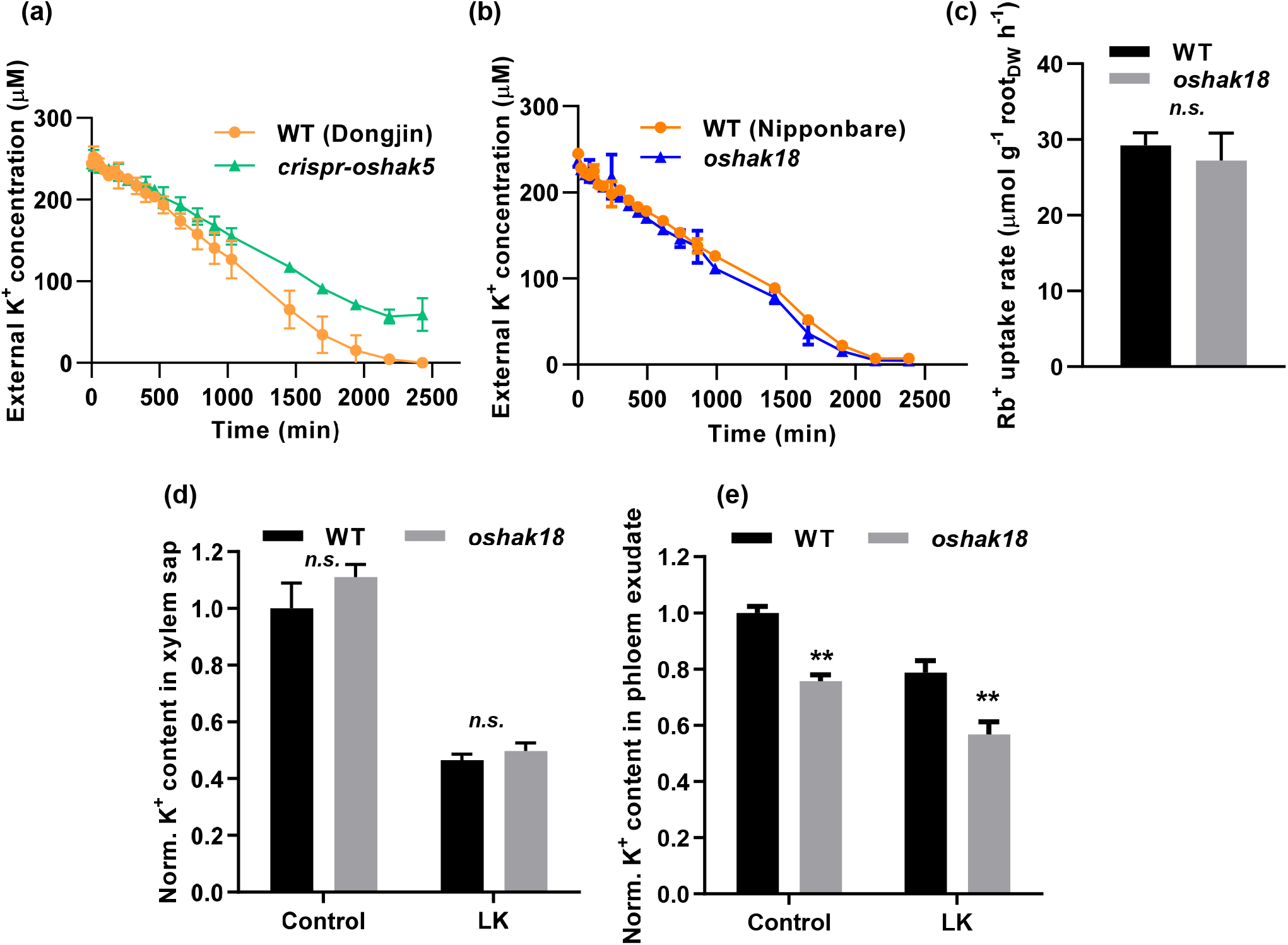
Disruption of *OsHAK18* doesn’t affect K^+^ uptake and xylem loading but inhibits phloem loading of K^+^. (a) Comparison of K^+^ uptake ability between WT and *oshak18* mutant. 7-d-old rice seedlings were pretreated in a solution without K^+^ at 28°C for 24 h, and then were transferred into a solution containing 250 μM K^+^ for K^+^ depletion assay. At each time point, 200 μL solution were sampled and determined for K^+^ concentration. (b) A CRISPR mutant of *OsHAK5* (*crispr-oshak5*) with a Dongjin background was used as a positive control in the K^+^ depletion assay. Data are shown as means ± SE (n = 3). (c) Comparison of Rb^+^ uptake rate between WT and *oshak18* mutant. 14 –d-old rice seedlings were starved of K^+^ for 14 days, and then were transferred to medium containing 50 μM Rb^+^ but no K^+^. After 6-h incubation, seedlings were harvested and determined for internal Rb^+^ contents. Rb^+^ absorption rates were calculated as described in previous study (Nieves-Cordones et al., 2019). (d) K^+^ concentrations in xylem sap collected from plants with or without LK treatment. Each data was normalized to the corresponding value of WT under HK control condition. Data bars are presented as means ± SD (n = 4). (e) K^+^ concentrations in phloem exudates. Each data was normalized to the corresponding value of WT under HK control condition. The data bars are shown as means ± SD (n = 4). Asterisks indicate significant differences from the WT (**, p < 0.01) by Student’s *t* test.

To reveal the function of OsHAK18 in K^+^ long-distance translocation, we measured the K+ concentration in the xylem sap and phloem sap collected from WT and *oshak18*. There were no substantial differences in xylem K^+^ concentration between WT and *oshak18* regardless of LK treatment (Fig. 7d). However, the K^+^ concentration in phloem exudates of *oshak18* was markedly lower than that of WT (Fig. 7e), suggesting that OsHAK18 plays crucial roles in K^+^ phloem loading and K^+^ redistribution.

### OsHAK18 affects the long-distance K^+^ redistribution from shoot to root

Split-root assays are usually carried out to assess the long-distance signaling of nutrient elements, such as Fe, S, P, and N (Ferreira Torres et al., 2021), which can also reflect the root-to-shoot-to-root translocation of nutrients. Thus, we conducted a split-root assay to investigate the effect of OsHAK18 on long-distance translocation of K^+^ in rice. After 9-d starvation for K^+^, each hydroponic-grown plant’s roots were physically divided into two similar isolated groups that remained attached to a common shoot. As shown in Fig. 8a, the two groups of roots were respectively immerged into HK or LK culture solution in separated zones of a special container. The K^+^ contents of roots in HK and LK zones were determined at 0, 1 and 3 day after root splitting. In HK zone, WT and *oshak18* had similar variation tendency on K^+^ contents from 0 to 3 day (Fig. 8b). The 1-d exposure in HK solution resulted in no significant changes on K^+^ contents of both WT and *oshak18*, but 3-d exposure dramatically increased the K+ contents (Fig. 8b), suggesting that there may be no differences on K^+^ uptake ability between WT and *oshak18*. In LK zone, the root K^+^ contents of WT and *oshak18* were still decreased with similar extent at 1 day compared with the corresponding values at 0 day (Fig. 8c), which may be due to the long-distance-induced delay of K^+^ translocation. After 3 days, the root K^+^ content of WT in LK zone increased intensively compared with the value at 1 day, which was also significantly higher than that of *oshak18* (Fig. 8c). This result indicates that the disruption of *OsHAK18* inhibits K^+^ long-distance translocation in rice. Given the localization and function of OsHAK18 in phloem tissue, we suggest that OsHAK18-mediated K^+^ phloem loading plays an important role in shoot-to-root redistribution of K^+^.

**Figure 8.**
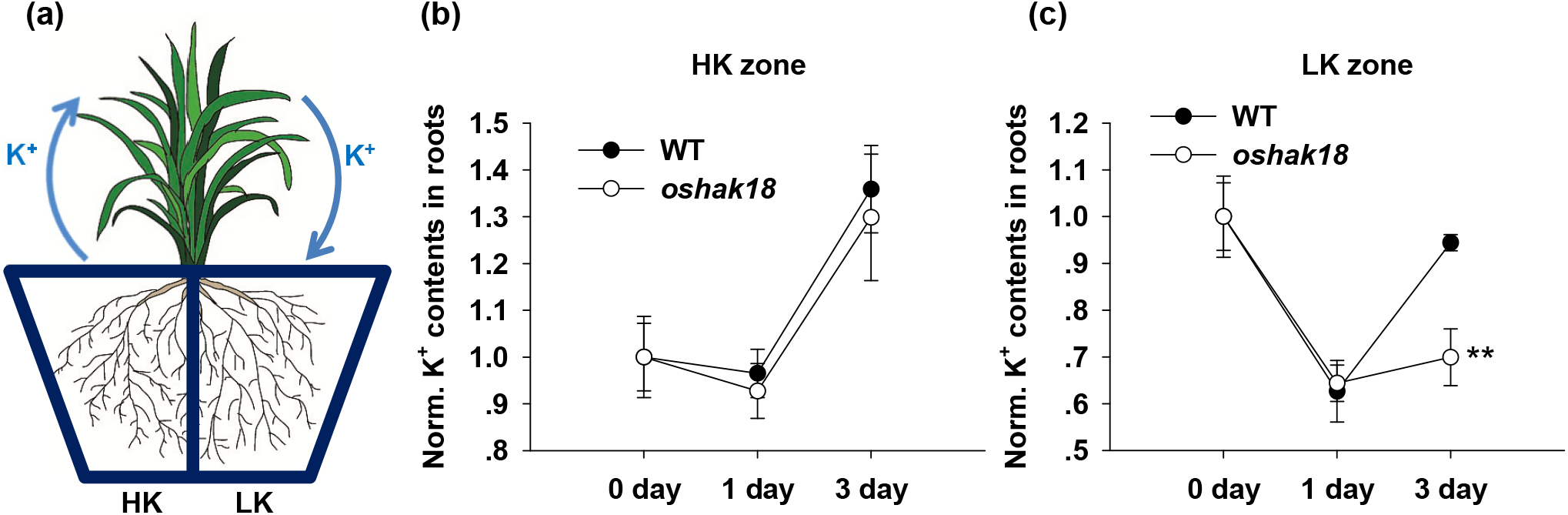
Split-root assay shows that disruption of *OsHAK18* inhibits the redistribution of K^+^. (a) Schematic diagram of the split-root system. 21-day-old rice seedlings were starved of K^+^ for additional 9 days and then were transferred to the special containers for split-root assay. The container was divided into two separate zones, which were filled with HK (5 mM) or LK (10 μM) hydroponic medium, respectively. The roots of rice seedlings were divided into two equal parts, which were immersed into the HK or LK culture solution for 3 days. At the indicated time points, the roots in HK or LK zone were sampled respectively and determined for K^+^ contents. The root K^+^ contents of WT or *oshak18* at each time point were normalized to their corresponding values at 0 day, which were set as standard 1. Data points represented means ± SD (n = 3). Asterisks indicate significant differences from the WT value at 3 day (**, p < 0.01) by Student’s *t* test.

### The effect of *OsHAK18* disruption on rice agronomic traits

We also compared the agronomic traits between WT and *oshak18* mutants at harvest stage. The rice plants were grown in the experimental fields outside with adequate water and fertilizer. Although there were no significant differences on plant height between WT and two CRISPR mutant lines (Fig. S7a,b), the effective tilling numbers of the two CRISPR mutant lines were significant lower than that of WT (Fig. S7c). By contrast, the grain number per panicle and seed setting percentage of the *oshak18* mutants were significantly higher than those of WT (Fig. S7d,e). However, no substantial differences were observed on the grain traits, including grain yield per plant, 100-grain weight and grain size (Fig. S7f-h).

## DISCUSSION

In low-K^+^ stressed Arabidopsis, the root K^+^ uptake was mainly mediated by the Shaker K^+^ channel AtAKT1 and AtHAK5 transporter (Pyo et al., 2010; Rubio et al., 2010). OsAKT1 has been demonstrated as the ortholog of AtAKT1 in rice, having similar function with AtAKT1 (Li et al., 2014). Interestingly, among the 13 KUP/HAK/KT transporters in Arabidopsis, AtHAK5 is the only one that is considered to play important roles in root K^+^ uptake. However, in rice, a few members in KUP/HAK/KT transporter family have been reported to be involved in root K^+^ uptake, such as OsHAK1, 5, 8, 16 (Chen et al., 2015; Feng et al., 2019; Wang et al., 2021b; Yang et al., 2014). Besides being expressed in root epidermis and cortex, these KUP/HAK/KT transporters are also present in vascular tissues, such as xylem parenchyma cells. Although they were suggested to mediate K^+^ release into xylem, the direct evidences are still absent. So their functions in vascular tissues remain unclear. In previous study, OsHAK21 was found expressed in passage cells in endodermis (Shen et al., 2015). Given the important roles of passage cells in nutrients’ entry into endodermis, OsHAK21 may also function in mediating root K^+^ uptake, especially when rice suffered salt stress (Shen et al., 2015). Recently, some KUP/HAK/KT transporters expressed in pollen, such as OsHAK1, 5, 26 and SlHAK5 (a tomato transporter) were shown to be essential to pollen viability (Li et al., 2022; Nieves-Cordones et al., 2020). Here, we reported OsHAK18, a phloem-localized KUP/HAK/KT potassium transporter, mediated phloem K^+^ loading to regulate K^+^ redistribution from shoot to root, which is a novel function of KUP/HAK/KT transporters. Taken together, the diversity of tissue expression of rice KUP/HAK/KT potassium transporters leads to the diversity of their functions.

We discovered that OsHAK18 mediated cell K^+^ absorption and showed no permeability to Na^+^ in yeast or *E. coli* (Fig. 1 and Fig. S2). According to the phylogenetic tree analysis in previous study (Very et al., 2014), OsHAK18 belongs to the subgroup III in KUP/HAK/KT family. Some members in this subgroup, such as OsHAK12 and AtKUP9, had been reported to perform the functions quite different from OsHAK18. In contrast with OsHAK18, OsHAK12 was shown to be not permeable to K^+^ but mediate Na^+^ uptake when expressed in yeast (Zhang et al., 2021). AtKUP9, an ER-localized KUP/HAK/KT transporter, was reported to mediate K^+^ and auxin efflux from ER to cytoplasm in root quiescent center (QC) cells to maintain meristem activity, which is essential to root growth under K^+^ deficiency (Zhang et al., 2020). Recently, Yamanashi et al. (2022) proposed that AtKUP9 participated in the K^+^ distribution in leaves and K^+^ uptake in roots under low K^+^ conditions (Yamanashi et al., 2022). The *atkup9* mutant accumulated more K^+^ in roots and has similar K^+^ level with WT in various above-ground parts (Yamanashi et al., 2022). Taken together these previous findings with our results here, we propose that OsHAK18 plays a different role form OsHAK12 and AtKUP9, although they have a close phylogenetic relationship. This may be due to the great differences on the subcellular localization, expression pattern or transporter characteristics among these transporters.

In rice, it has been revealed that the Shaker K^+^ channel, OsAKT2, is a key player mediating phloem K^+^ loading. Disruption of *OsAKT2* resulted in a significant decrease of K^+^ concentration in phloem sap, and restrained K^+^ redistribution (Tian et al., 2021). In this study, we found that OsHAK18, similar to OsAKT2, also played important roles in phloem K^+^ loading. Compared with WT, the *oshak18* mutants had lower K^+^ concentration in phloem sap and a reduced K^+^ redistribution (Fig. 7e and 8c). However, it should be noted that the effects of these two K^+^ transporters on grain traits are quite different. Disruption of *OsAKT2* results in slender grains and a significant reduction of grain weight and yield, whereas there are no substantial differences on grain traits between *oshak18* mutants and WT (Fig. S7h-g). It had been revealed that enhanced sucrose loading into phloem can improve rice grain yield by increasing grain size (Wang et al., 2015), which is mainly mediated by sucrose/H^+^ symporters (SUT/SUCs). However, SUT/SUC transporters need to be driven by transmembrane electrochemical gradient or hyperpolarized membrane potential (Dreyer et al., 2017). The AKT2-like K^+^ channels can behave as ‘open-lock’ channels mediating K^+^ efflux, maintaining a relative hyperpolarized membrane potential to promotes SUT/SUC-mediated sucrose loading (Dreyer et al., 2017; Gajdanowicz et al., 2011). Hence, OsAKT2 was suggested to control the grain size and yield by facilitating sucrose phloem loading and long-distance translocation (Shen et al., 2020; Tian et al., 2021). However, although OsHAK18 functions in mediating phloem K^+^ loading too, it is not likely to participate in maintaining membrane potential and facilitate sucrose phloem loading, as what the AKT2-like channels do. Hence, no differences on grain traits were detected between WT and *oshak18* mutants.

In this study, we showed that OsHAK18 operated as a typical KUP/HAK/KT transporter mediating cell K^+^ absorption and was preferentially expressed in phloem tissue (Fig. 2). Interestingly, disruption of *OsHAK18* rendered rice seedlings insensitive to low-K^+^ stress (Fig. 3). Compared with WT, the *oshak18* mutants accumulated more K^+^ in shoots but less K^+^ in roots (Fig. 4,5,6). OsHAK18 disruption also significantly improved the shoot/root ratios of K^+^ per plant and decreased the K^+^ level in phloem sap (Fig. 4 and Fig. 7). We proposed that disruption of *OsHAK18* facilitate K^+^ retention in shoots, thereby rendering rice insensitive to low-K^+^ stress. However, because OsHAK18 doesn’t affect root K^+^ uptake (Fig. 7b,c), mutating this gene alone is not likely to generate durable resistance to low-K^+^ stress. Therefore, if OsHAK18 disruption is combined with some strategies of improving rice K^+^ absorption, such as increasing OsHAK1, 5 or OsAKT1activities, it will be promising to improve rice durable tolerance to K^+^ deficiency.

## MATERIALS AND METHODS

### Functional characterization of OsHAK18 in yeast and *E. coli* and Arabidopsis *athak5*

The coding sequence of *OsHAK18* with the locus number Os09g0563200 (or LOC_Os09g38960) was obtained from the Rice Genome Annotation Project (http://rice.uga.edu/). For *OsHAK18* expression in yeast, *E. coli* and *Arabidopsis athak5* mutant, the coding sequence of *OsHAK18* was subcloned into a yeast expression vector (pYES2), an *E. coli* expression vector (pEASY-Blunt), and a binary vector (pCAMBIA1307), respectively. The primers used were listed in Table S1.

The recombinant vector *pYES2-OsHAK18* or empty *pYES2* were transformed into the K^+^ uptake-deficient strain CY162 (*MATα, Δtrk1, trk2::pCK64, his3, leu2, ura3, trp1 and ade2*) using the LiAC/ssDNA/PEG method, and the growth assays on arginine phosphate (AP) medium containing different concentrations of K^+^ were performed as described previously (Horie et al., 2011a; Tian et al., 2021).

For functional analysis of OsHAK18 in *E. coli, pEASY-OsHAK18* or the empty vector were transformed into the K^+^ uptake-deficient *E. coli* stain, TK2420 (a triple mutant that is defective for the Kdp, Trk and Kup K^+^ uptake systems). The positive transformants were cultured in the LBK medium containing 10 g/L tryptone, 5 g/L yeast extract, 87 mM KCl and 0.1 g/L ampicillin till the absorbance (*A*_600 nm_) reached 0.8-1.0. After washed three times with deionized water, the transformants were suspended to 1.0 (*A*_600 nm_) and dripped on the minimal medium (MM) containing 0.5 mM IPTG and different K^+^ concentrations, which was described in previous study (Cheng et al., 2016). To plot the cell growth curve, the transformants were diluted to 0.01-0.02 after 4 h-incubation of 0.5 mM IPTG, and then were cultured in MM medium with 0.5 mM IPTG, 0.1 g/L ampicillin and different K^+^ concentrations.

To verify the K^+^ transporter activity of OsHAK18 in *athak5*, the OsHAK18 coding sequence was amplified and built into the pCAMBIA1307 vector driven by 2×35S promoter. The recombinant plasmid was introduced into *Ag. tumefaciens* strain GV3101 to infect *athak5* inflorescences using the floral dipping method. The homozygous transgenic lines of the T3 generation were used for phenotype analysis under LK conditions, which was performed as described previously (Pyo et al., 2010).

### Subcellular localization and GUS staining assay

The *OsHAK18* coding sequence without the stop codon was coloned into the pUC-EGFP vector. The recombinant plasmid was coated with 0.6 μm gold particles (1 μg DNA per 10 mg gold) and then introduced into onion epidermal cells using the particle bombardment method (PDS-1000/He particle delivery system, Bio-Rad). The epidermis was incubated on MS medium in darkness at room temperature (20–25 °C) overnight before fluorescence analysis using confocal laser microscopy (LSM780, Carl Zeiss). The coding sequence of *OsHAK18* without the stop codon was fused with EGFP at the C-terminal in the pSuper1300 vector. The recombinant plasmids, pSuper1300-OsHAK18:EGFP, were transformed into *Agrobacterium* strain GV3101. OsAKT1:mCherry was used as a marker to label plasma membrane, which was co-expressed with OsHAK18:EGFP. Tobacco leaves were transfected using the *Agrobacterium* infiltration method and incubated for 2 days before fluorescence assay.

The promoter of *OsHAK18* (a 2637-bp fragment upstream from the ATG start codon) was cloned from genomic DNA using the primer set shown in Table S1, and then constructed into the pCAMBIA1301 vector to replace the region containing LacZ and 35S promoter. The recombinant plasmid was introduced into *Ag. tumefaciens* strain EHA105 and transformed into rice calli as previously described (Nishimura et al., 2006). The T2 heterozygous transgenic rice plants were used for GUS histochemical staining which was performed as described previously (Shen et al., 2020).

### Plant materials and growth conditions

The *oshak18* mutant line (accession number: NE5666) with a Nipponbare background was obtained from the rice *Tos17* insertion mutant database (https://tos.nias.affrc.go.jp) (Miyao et al., 2003). Homozygous mutant plants were identified by PCR using the gene-specific primers listed in Table S1. Rice plants (*Oryza sativa* L. ssp. japonica cv. Nipponbare) were used as wild-type (WT) controls unless otherwise stated.

To generate *OsHAK18* knockout mutants using the CRISPR/Cas9 editing system, we designed two specific guide RNA (gRNA) sequences using CRISPR-PLANT database (https://www.genome.arizona.edu/crispr), which was listed in Table S1. The two gRNA spacers were cloned into the *Aar I* position of the SK-gRNA vector, respectively, following a method described previously (Shen et al., 2017). These intermediate vectors were digested by *Bgl* II and *Kpn* I, generating the cassettes which were put into the *BamH* I and *Kpn* I position of pC1300-Cas9. Then the recombinant vectors were transformed into callus of WT (Nipponbare) via *Agrobacterium* EHA105 to generate CRISPR mutant lines. The mutations of *OsHAK18* were confirmed by DNA sequencing in T2 generation of the two independent lines (Fig. S3c). The T3 progeny was used in phenotypic assay and K^+^ content determination.

To construct *OsHAK18* overexpression lines, pSuper1300-OsHAK18:EGFP was transformed into the callus of WT (Nipponbare) via *Agrobacterium* EHA105. The homozygous T3 generation lines were applied to determine the expression level and K^+^ contents.

For phenotypic analysis, rice seeds were sterilized in 2% (v/v) sodium hypochlorite for 1 h, rinsed thoroughly with deionized water and immersed in water for 2 d at 37 °C in darkness. Germinated seeds with similar sprouts were selected for hydroponic cultivation and cultured in a phytotron with a 14-h-light (28°C), 10-h-dark (24°C) photoperiod. After grown in water for 7 d, the seedlings were deprived of endosperms to avoid possible involvement of K^+^ supplied from this organ. Then the seedlings were transferred to the International Rice Research Institute (IRRI) nutrient solution (Yoshida et al., 1971), and cultured for additional 7 d. The pH was adjusted to 5.8 with KOH (final K^+^ concentration was approximately 3.5 mM). For LK treatment, 14-day-old rice seedlings were transferred into low-K^+^ (LK) IRRI medium, in which K_2_SO_4_ was substituted by equal concentration of Na_2_SO_4_. The pH of LK medium was adjusted to 5.8 with NaOH. The hydroponic medium was changed for fresh every 3 d.

### Kinetic analysis of K^+^ uptake and Rb^+^ uptake rate determination

To compare root K^+^ uptake activity among different genotypes, we performed K^+^-depletion experiments as described in previous study (Li et al., 2014) with some modification. 7-d-old seedlings were pretreated in a K^+^-free solution (0.2 mM CaSO_4_ and 5 mM MES, pH 5.8 adjusted with Tris) at 28°C for 24 h. Each sample included seven seedlings, whose weight was approximately 0.8 g. After a 24-h K^+^ starvation, the samples were transferred to the depletion solution (0.25 mM KNO_3_, 0.2 mM CaSO_4_, and 5 mM MES, pH 5.8 adjusted with Tris). The samples were put on a shaking table at 28°C in a 14-h-light/10-h-dark photoperiod. At each time point, 200 μL solution were sampled and determined for K^+^ concentration.

For Rb^+^ uptake rate determination, 14–d-old rice seedlings were treated with LK medium for additional 12 d, and then the K^+^-starved seedlings were transferred to IRRI solution containing 50 μM Rb^+^ but no K^+^. After 6-h incubation, seedlings were harvested and determined for internal Rb^+^ contents. Rb^+^ absorption rates were calculated as described previously (Nieves-Cordones et al., 2019b).

### Determination of ion content in xylem sap and phloem exudate

The xylem sap and phloem exudate were collected as described in previous study (Tian et al., 2021) with some modification. Briefly, the hydroponically cultured 14-day-old rice seedlings were transferred to LK medium. After 9-d treatment, the xylem sap and phloem exudate were collected, respectively. Shoots were cut at about 2–3 cm above the junction between roots and shoots, and then a cotton ball was attached to the cut surface of stump, which was then covered by an inverted centrifuge tube. The collection lasted for 3 hr. Then the xylem sap was squeezed from the cotton balls by centrifugation method. To collect phloem exudate, the cut surface of shoots excised from seedlings was immersed into a 15 mM Na_2-_EDTA buffer (pH 7.5). To avoid blockage of the cut surface, the endmost parts were consecutively recut at 1 mm from the end three times at 1-min intervals in the EDTA solution. Then the shoots were transferred into the EDTA solution and incubated in a dark chamber for 8 hr at 90% humidity. The K^+^ concentration in the xylem sap and phloem exudate were determined using inductively coupled plasma/optical emission spectrometry (PerkinElmer, Waltham, USA).

### Split-root assay

21-day-old rice seedlings were transferred to LK hydroponic medium and pretreated for 10 days before split-root assays. Then, the roots of each plant were divided into two groups by hand, and each group was submerged into hydroponic medium containing 5 mM (HK) or 10 μM K^+^ (LK), respectively. The HK and LK medium were filled into two different zones of a container, which were separated by a central partition. The junction of shoots and roots were wrapped with sponge and fixed in holes on polystyrene foam plates. At each time point, the roots in HK or LK zones were sampled respectively, and were determined for K^+^ content.

## ACKNOWLEDGEMENTS

This work was supported by grants from the National Natural Science Foundation of China (No. 32270268 and No. 32171956); the open funds of the State Key Laboratory of Plant Physiology and Biochemistry (No. SKLPPBKF1904); the Natural Science Foundation of Jiangsu province in China (No. BK20140699).

## AUTHOR CONTRIBUTIONS

L.S. conceived and designed the research; L.S., Q.W., W.F, J.L., N.L., D.C, Q.T. and W.J. performed the experiments. L.S. and Q.W. analyzed the data. W.Z supervised the laboratory work. L.S. wrote and revised the manuscript.

## DATA AVAILABILITY STATEMENT

The data that support the findings of this study are available from the corresponding author upon reasonable request.

