## supporting information for "Rice potassium transporter OsHAK18 mediates phloem K^+^ loading and redistribution"

**Supplementary Figure 1**

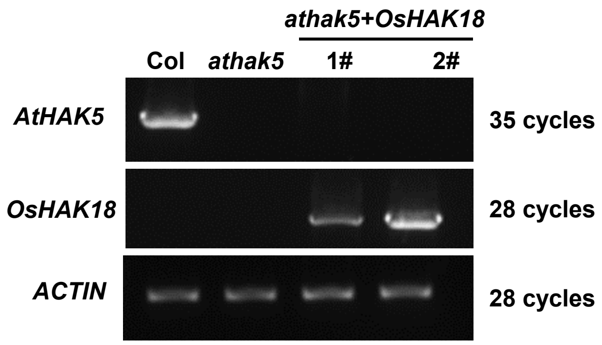


**Figure S1. Identification of *OsHAK18* expression in transgenic *Arabidopsis*.** RT-PCR analysis using cDNA derived from wild-type (Col), *athak5* and two independent lines of *OsHAK18*-expressed *athak5* (*athak5*+*OsHAK18*, 1# and 2#). *ACTIN* was used as an internal control.

**Supplementary Figure 2**

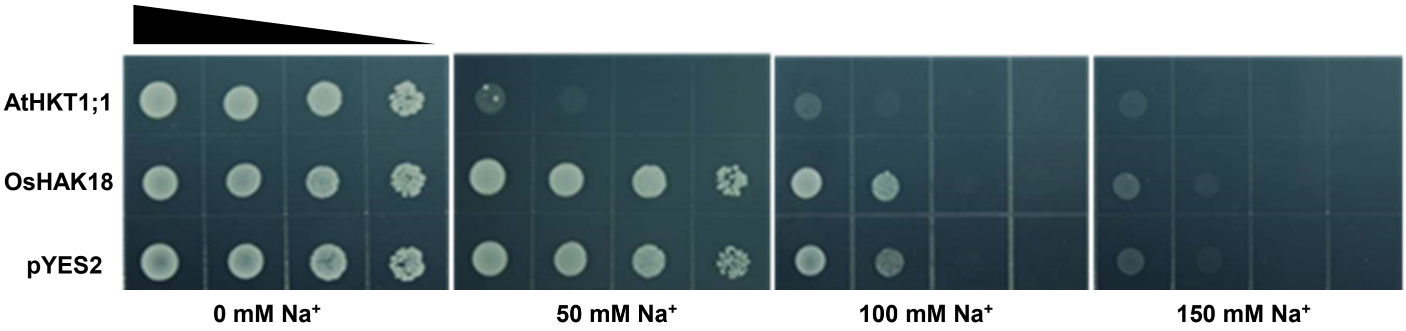


**Figure S2. Functional analysis of OsHAK18 in $Na^+$-sensitive yeast (B31).** Growth status of $Na^+$-sensitive yeast (B31) expressing *OsHAK18*, *AtHKT1;1* or empty vector (pYES2) on arginine phosphate (AP) medium supplemented with different concentrations of NaCl (0, 50, 100, 150 mM). Black triangle indicates 1:10 serial dilutions of yeast cells placed on the mediums.

Supplementary Figure 3

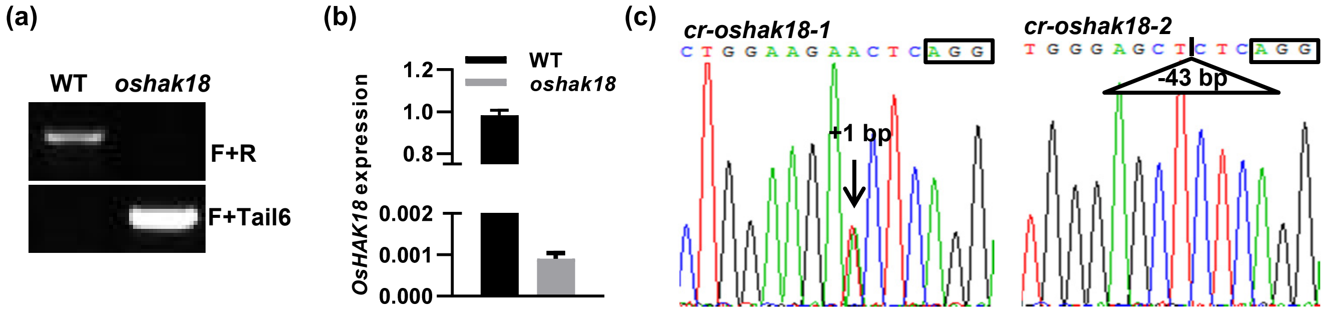


**Figure S3. Identification of *OsHAK18 Tos17* insertion mutant and CRISPR lines.** (a) Identification of the *Tos17* insertion and the homologous *oshak18* mutant. Genomic DNA extracted from WT and *oshak18* was used as the PCR templates, and the specific primer sequences were listed in Table S1. F and R represent the primers located in the flanking sequences of the *Tos17* insertion site, while Tail6 is a primer located at the 3′ end of *Tos17.* (b) Comparison of *OsHAK18* expression level between WT and *oshak18* using qRT-PCR analysis. *OsUBQ5* was used as an internal control. (c) The effect of OsHAKe8 CRISPR was verified by amplicon sequencing in *cr-oshak18-1* and *cr-oshak18-2*. The arrow indicates one base (A or T) insertion in *cr-oshak18-1*, while the triangle represents a 43-bp deletion in *cr-oshak18-2* The black boxes show the PAM sites.

**Supplementary Figure 4**

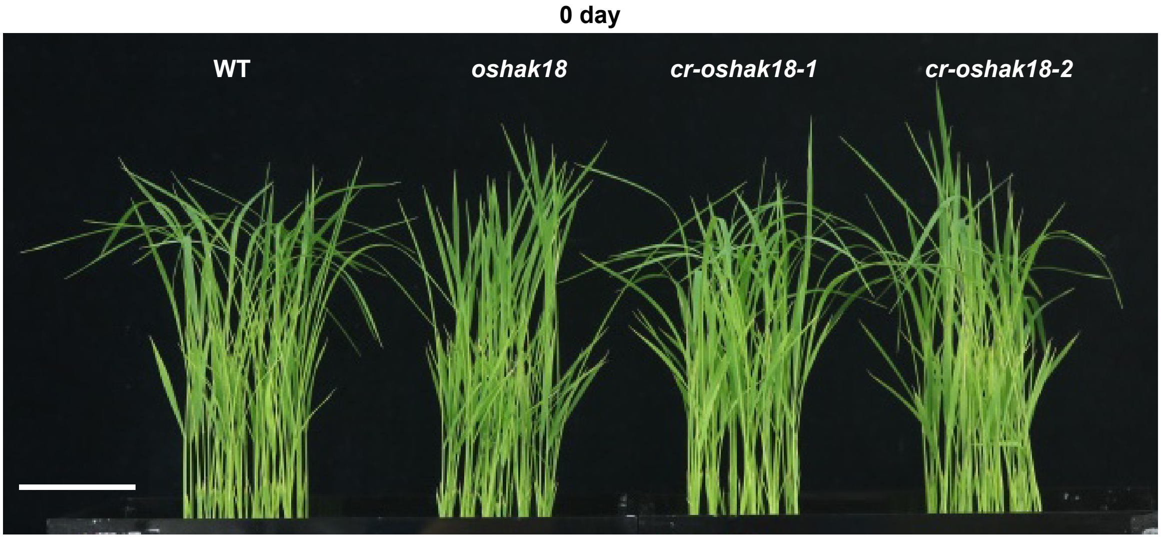


**Figure S4. Phenotypic comparison among different genotypes before LK treatment.** Two-week-old seedlings were photographed before LK treatment (o day). Bar = 5 cm

**Supplementary Figure 5**

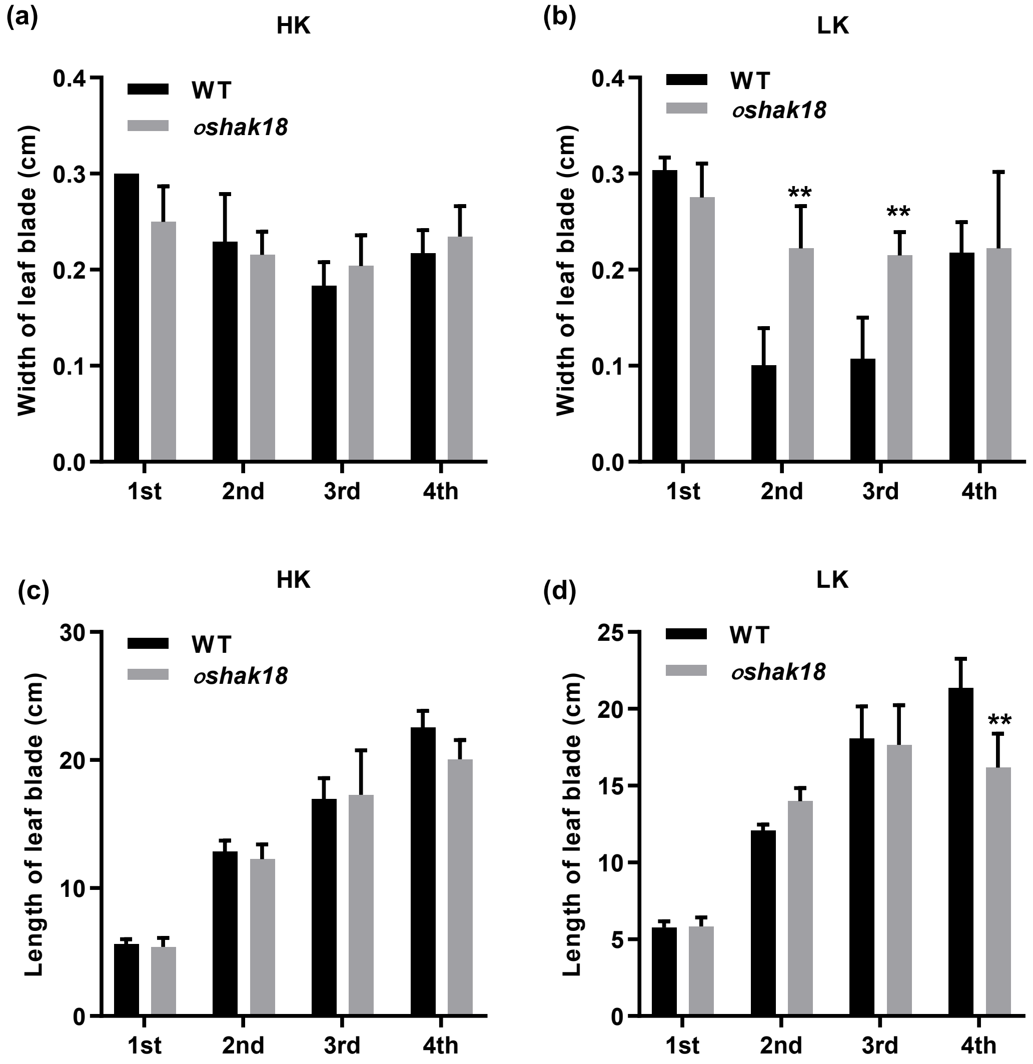


**Figure S5. The width and length of leaf blades.** Two-week-old seedlings were transferred to low-K+ (LK) hydroponic medium containing 10 μM K+, and treated for 12 days. The numbers 1st to 4th on X-axis represent the four leaves numbered in order of development. (a,b) The width of leaves under control and LK condition. (c,d) The length of leaves under control and LK condition. ). Each data bar was shown as means $\pm$ SD (n = 10). Asterisks indicate significant differences from the WT (**, $p < 0.01$) by Student's *t* test.

Supplementary Figure 6

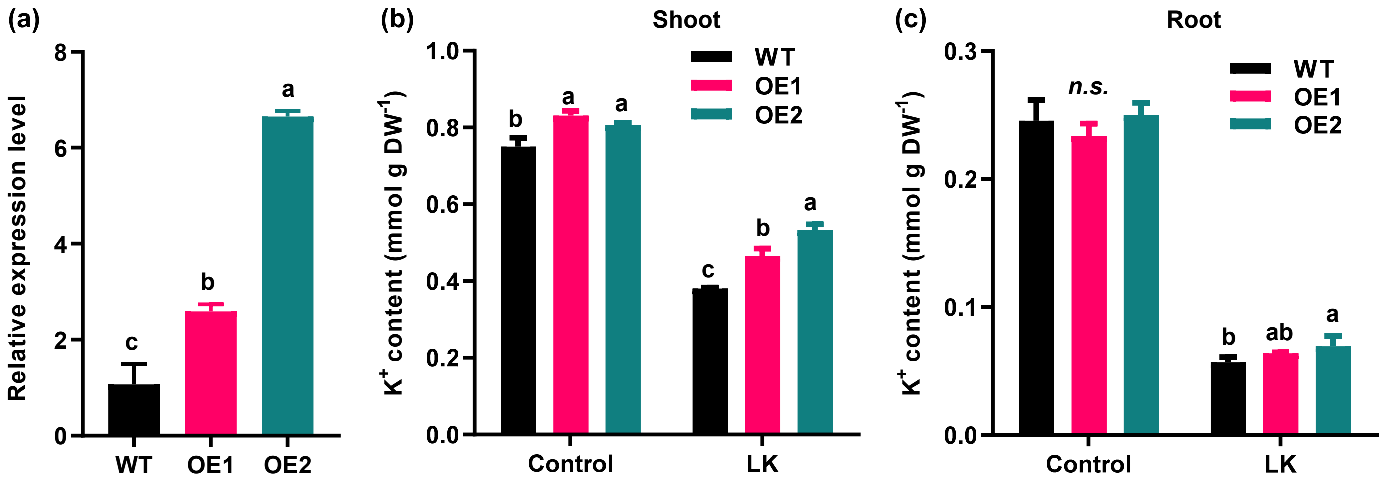


**Figure S6. Comparison of K⁺ content between WT and *OsHAK18*-overexpression lines.** (a) Identification of *OsHAK18*-overexpression lines (OE1 and OE2) using qRT-PCR. (b,c) K+ contents in shoots and roots. Two-week-old seedlings were transferred to control (containing 5 mM $K^+$) or LK medium (containing 10 μM $K^+$), and treated for 12 days. (b,c) $K^+$ contents in shoots and roots. Each data bar was shown as means ± SD (n ≥ 3). Statistical significance analyzes were carried out by one-way analysis of variance (ANOVA) with Tukey's multiple range test at $P < 0.05$.

### Supplementary Figure 7

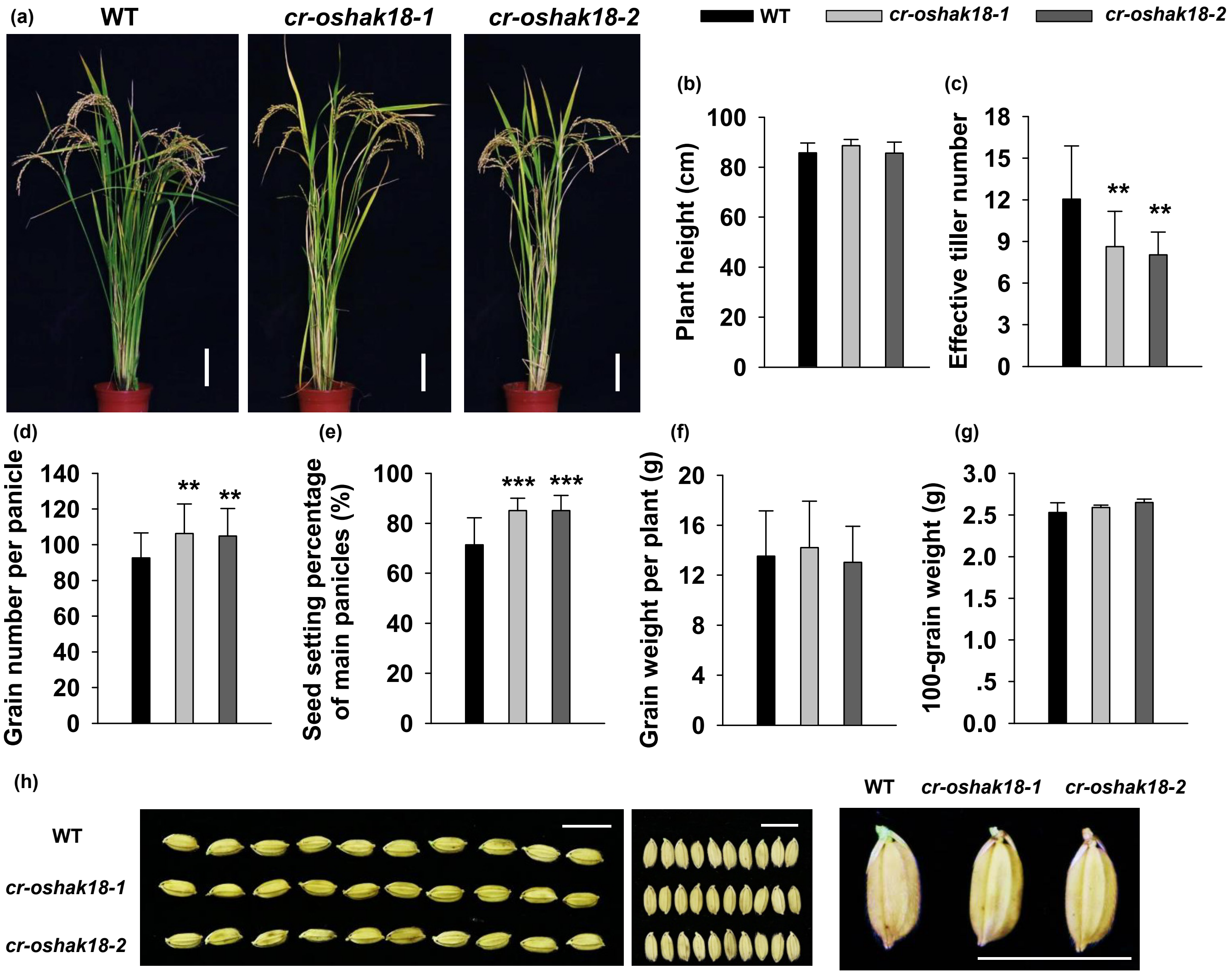


**Figure S7. The comparison of agronomic traits between WT and *oshak18* CRISPR lines under the field experiment conditions.** (a) Gross morphology comparison between WT and *oshak18* CRISPR lines at harvest stage. Bar = 10 cm. (b) Plant length (n ≥ 17). (c) Effective tiller number (n ≥ 17). (d) Grain number per panicle (n ≥ 40). (e) Seed-setting percentage of main panicles (n ≥ 40). (f) Grain weight per plant (n ≥ 30). (g) 100-grain weight (n = 9). (h) Comparison of grains from WT and *oshak18* CRISPR lines. Bar = 10 mm. The data bars in (b) to (g) represent the means ± SD. Asterisks indicate significant differences from the WT (**, $P < 0.01$; ***, $P < 0.001$) by Student's *t*-test.

**Supplementary Table 1**

**TABLE S1. PCR Primers used in this study**

| Name | Sequence (5' to 3') | Purpose/vector |
|---|---|---|
| pYES2-F | cccaagcttATGGAGACCAGAACAAATGAGT | pYES2 |
| pYES2-R | tgctctagaTTACACGTAGAAAACCTGCC | pYES2 |
| pEASY-F | ATGGAGACCAGAACAAATGAGT | pEASY |
| pEASY-R | CACGTAGAAAACCTGCCCAA | pEASY |
| 1307-F | cgctctagaactagtggatccATGGAGACCAGAACAAATGAGT | pCAMBIA1307-Flag |
| 1307-R | atcatggtctttgtagtcgacCACGTAGAAAACCTGCCCAA | pCAMBIA1307-Flag |
| AtHAK5-F | ATGGATGGTGAGGAACATCAAATAG | RT-PCR |
| AtHAK5-R | TTATAACTCATAGGTCATGCCAACC | RT-PCR |
| ACTIN-F | CTCAGGTATTGCAGACCGTATGAG | RT-PCR |
| ACTIN-R | CTGGACCTGCTTCATCATACTCTG | RT-PCR |
| pUC-EGFP-F | tgctctagaATGGAGACCAGAACAAATGAGTA | Localization in onion epidermis |
| pUC-EGFP-R | tcccccgggCACGTAGAAAACCTGCCCAA | Localization in onion epidermis |
| 1300GFP-F | cccaagcttATGGAGACCAGAACAAATGAGTA | Localization in tobacco and OE |
| 1300GFP-R | cggggtaccCACGTAGAAAACCTGCCCAA | Localization in tobacco and OE |
| ProOsHAK18-F | tgctctagaTGGATTTGAGGTGATATGCTGG | GUS/pCAMBIA1301 |
| ProOsHAK18-R | catgccatggGGGTTCAGACTTCAGATCAATCAAC | GUS/pCAMBIA1301 |
| F | TTTCCCAAGTCGGTCTGAATGATAC | Identification of Tos17 insertion |
| R | CGTATATCATCTCTCCACGCAGAAC | Identification of Tos17 insertion |
| Tail6 | AGGTTGCAAGTTAGTTAAGA | Identification of Tos17 insertion |
| qRT-F | ACAATGGCCAAGGAGGAACA | qRT-PCR for OsHAK18 |
| qRT-R | TTCATATGTGCGGCGACTGT | qRT-PCR for OsHAK18 |
| OsUBQ5-F | ACCACTTCGACCGCCACTACT | qRT-PCR |
| OsUBQ5-R | ACGCCTAAGCCTGCTGGTT | qRT-PCR |
| CRISPR-T1-F | ggcaATGCAGAGGCTGGAAGACTC | CRISPR/SK-gRNA |
| CRISPR-T1-R | aaacGAGTCTTCCAGCCTCTGCAT | CRISPR/SK-gRNA |
| CRISPR-T2-F | ggcaGGGCAATGTGGGAGCTGGAA | CRISPR/SK-gRNA |
| CRISPR-T2-R | aaacTTCCAGCTCCCACATTGCCC | CRISPR/SK-gRNA |
| Seq-F | CCTGGTATTCTGATTGGTCCTTTTA | Sequencing CRISPR mutataion |
| Seq-R | ATTTACTCCCTTACACTCATTCCAC | Sequencing CRISPR mutataion |
